# Comprehensive analysis of epigenetic signatures of human transcription control^†^

**DOI:** 10.1101/2020.09.23.309625

**Authors:** Guillaume Devailly, Anagha Joshi

**Affiliations:** GenPhySE, Université de Toulouse, INRAE, ENVT, 31326, Castanet Tolosan, France; Computational Biology Unit, Department of Clinical Science, University of Bergen, 5021, Bergen, Norway.

## Abstract

Advances in sequencing technologies have enabled exploration of epigenetic and transcription profiles at a genome-wide level. The epigenetic and transcriptional landscape is now available in hundreds of mammalian cell and tissue contexts. Many studies have performed multi-omics analyses using these datasets to enhance our understanding of relationships between epigenetic modifications and transcription regulation. Nevertheless, most studies so far have focused on the promoters/enhancers and transcription start sites, and other features of transcription control including exons, introns and transcription termination remain under explored. We investigated interplay between epigenetic modifications and diverse transcription features using the data generated by the Roadmap Epigenomics project. A comprehensive analysis of histone modifications, DNA methylation, and RNA-seq data of about thirty human cell lines and tissue types, allowed us to confirm the generality of previously described relations, as well as to generate new hypotheses about the interplay between epigenetic modifications and transcript features. Importantly, our analysis included previously under-explored features of transcription control namely, transcription termination sites, exon-intron boundaries, middle exons and exon inclusion ratio. We have made the analyses freely available to the scientific community at joshiapps.cbu.uib.no/perepigenomics_app/ for easy exploration, validation and hypotheses generation.

## Background

Epigenetic modifications of the DNA sequence and DNA-associated proteins along with transcriptional machinery is thought to be the main driver shaping mammalian genome during development and disease ^1^. Epigenetic modifications include DNA methylation, histone variants and histone post-translational modifications (such as acetylations and methylations), and facilitate tissue specific expression ^2^. The advent and maturation of sequencing technologies have facilitated large scale generation of epigenomic data across diverse organisms in multiple cell and tissue types. Accordingly, consortia were established to generate large epigenomic datasets, including ENCODE ^3^, Roadmap Epigenomics ^2^, Blueprint epigenome ^4^ for human, modENCODE ^5^ for model organisms, and FAANG ^6,7^ for farm animal species. The International Human Epigenome Consortium (IHEC) was set up to gather reference maps of human epigenomes ^8^. The data from these efforts have generated new findings through integrated analyses. Such analyses are facilitated by consortia data portals ^8,9^ as well as portals gathering data from multiple sources ^10–14^, which allow easy browsing as well as downloading both sequences and processed data. In addition, many data portals include (or link to) genome browsers to allow online solutions for the data exploration ^15^.

Several online tools have been developed to explore publicly available epigenomic data ^16–20^ to gain insights into mammalian epigenetic control. These tools used diverse computational frameworks ranging from data integration and visualisation (e.g. ChIP-Atlas allows visualisation multiple histone modifications and transcription factor binding sites at given genomic locus by using public ChIP-seq and DNase-seq data ^21^) to semi-automated annotation of genome (e.g. Segway performed genomic segmentation of human chromatin by integrating histone modifications, transcription-factor binding and open chromatin ^22^). Though identification of functional elements from epigenetic data ^2,22^ has been highly effective to annotate the enhancer and promoter regions of a genome, they failed to capture other transcription regulation features such as exon-intron boundaries and transcription termination features. For example, Roadmap Epigenomics project mapped about 30 epigenetic modifications across human cell lines and tissues to gather a representative set of “complete” epigenomes ^2^. Using this data, Kundaje *et al*. built a hidden Markov model based classifier to define 15 distinct chromatin states, including active or inactive promoters, active or inactive enhancers, condense and quiescent states. Notably, this unsupervised approach did not lead to the definition of “exon” states, and even less to “exon-included” and “exon-excluded” states. This might be because epigenetic modifications enriched at enhancers and promoters have a strong signal (or peaks), while the ones abundant at gene bodies (DNA methylation, H3K36me3) are wide and diffuse. The promoter and enhancers features therefore dominate in epigenetic data analyses, hindering recovery of associations between epigenetic modifications and other transcription control events such as splicing (constitutive or alternative). Moreover, some transcription features might not have a strong correlation with any chromatin modification studied. For example, Curado *et al*. ^23^ have estimated that only 4% of differential included exons were associated with changes in H3K9ac, H3K27ac, and/or H3K4me3 across 5 different cell lines. A gap therefore remains in genome-wide computational analyses towards getting a comprehensive overview of the associations between epigenetic modifications and transcription control features.

On the other hand, individual targeted studies have provided evidence for interplay between epigenetics and other transcriptional features. DNA methylation at gene bodies has been positively correlated with gene expression level ^24,25^. Maunakea *et al.* ^26^ observed that DNA methylation was positively correlated with splicing at alternatively spliced exons, and proposed a mechanism involving DNA methylation reader MECP2. Lev Maor*et al*. ^27^ observed that DNA methylation at exons can be either positively or negatively correlated with splicing depending on the exons, through a mechanism involving CTCF and MECP2. A causal role of DNA methylation in alternative splicing was established by drug-induced de-menthylation ^28^, as well as by targeted DNA methylations and de-methylations by Shayevitch *et al*. ^29^. Xu *et al*. ^30,31^ identified H3K36me3 epigenetic modification associated with alternative splicing. There is some evidence for epigenetic control at transcription termination as well: the loss of gene body DNA methylation was found to favour the usage of a proximal alternative poly-adenylation site by unmasking CTCF binding sites ^32^.

In summary, many large data integration approaches only allow extraction of epigenetic signatures for dominant features of transcription control (e.g. enhancer, promoter, TSS), missing many other transcription features (e.g. exon, intron, TSS). We therefore performed a systematic analysis of associations between epigenetic modifications and diverse transcription features. Using the Roadmap Epigenomics project data for 30 epigenetic modifications in about 30 cell and tissue contexts, we explored links between epigenetic modifications and transcription control. We confirmed previously known associations as well as generated novel observations. We have provided our analyses freely through a web application available at https://joshiapps.cbu.uib.no/perepigenomics_app/, allowing researchers to browse the results and generate working hypotheses.

## Results

### Exploration of epigenetic signatures of transcription control using the Roadmap Epigenomics project data

To explore the epigenetic signatures at transcription control sites, we first extracted three gene features: Transcription start site (TSS), transcription termination site (TTS) and middle exons from GENCODE annotation version 29^33^ (Table 1). We classified genes in two different ways. Firstly, we partitioned all genes based on the gene length “long” (*>*3kb), “short” (*≤*1kb), and “intermediate” length genes. We also classified genes based on a simplified GENCODE gene types, namely: protein coding, RNA genes, pseudogenes and other genes (see method section).

**Table 1.**
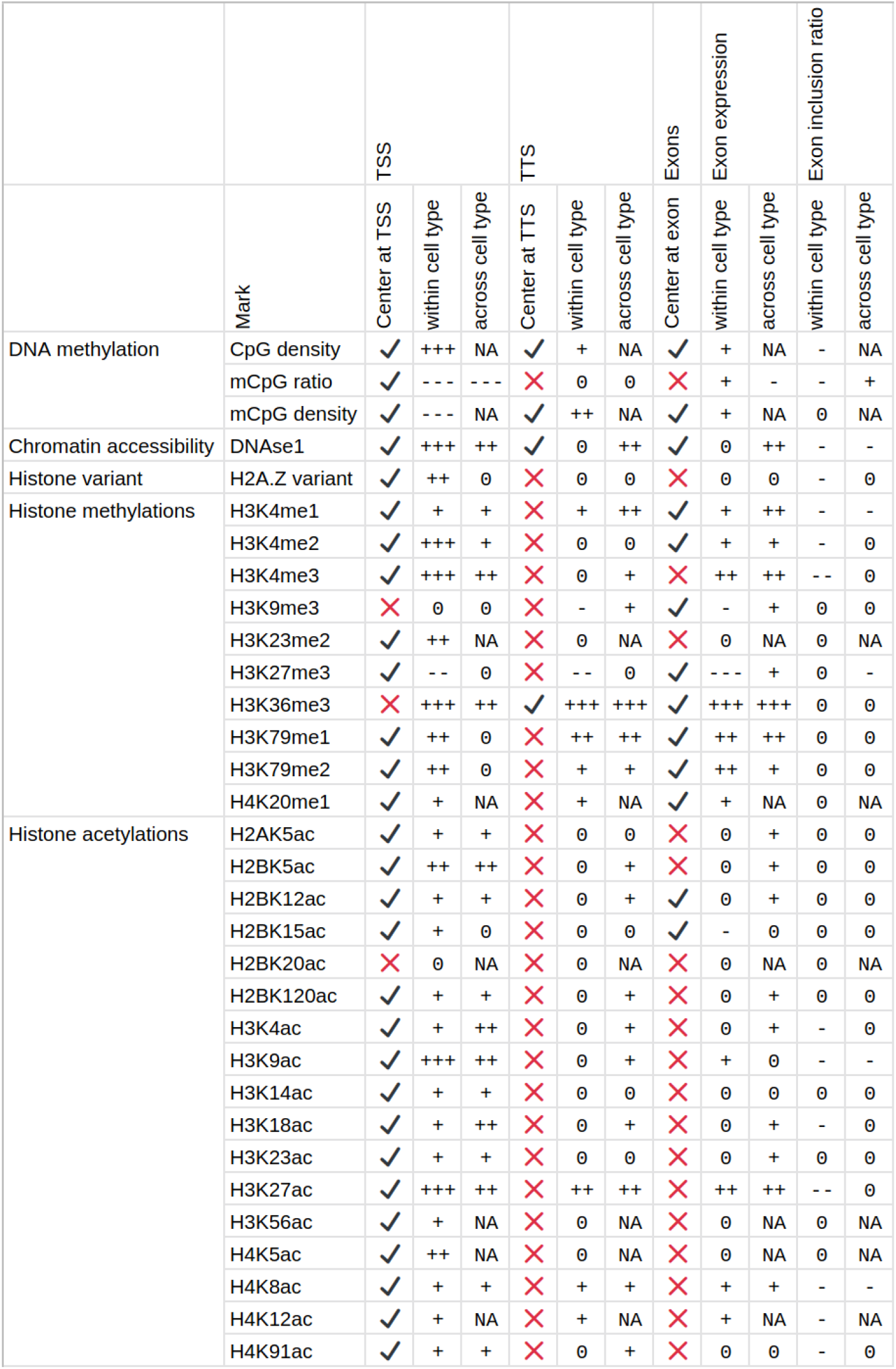
Summary of the associations between epigenetic modifications and transcription features in the Roadmap epigenomics project data. Tick (or cross): whether (or not) an epigenetic assay shows an increase or decrease in signal at the feature. Pluses: positive associations. Minuses: negative associations. Zeroes: no correlation. NA: not available (data not available in enough cell types).

|  |  | TSS |  |  | TTS |  |  | Exons | Exon expression |  | Exon inclusion ratio |  |
| --- | --- | --- | --- | --- | --- | --- | --- | --- | --- | --- | --- | --- |
|  | Mark | Center at TSS | within cell type | across cell type | Center at TTS | within cell type | across cell type | Center at exon | within cell type | across cell type | within cell type | across cell type |
| DNA methylation | CpG density | ✓ | +++ | NA | ✓ | + | NA | ✓ | + | NA | - | NA |
|  | mCpG ratio | ✓ | --- | --- | ✗ | 0 | 0 | ✗ | + | - | - | + |
|  | mCpG density | ✓ | --- | NA | ✓ | ++ | NA | ✓ | + | NA | 0 | NA |
| Chromatin accessibility | DNAse1 | ✓ | +++ | ++ | ✓ | 0 | ++ | ✓ | 0 | ++ | - | - |
| Histone variant | H2A.Z variant | ✓ | ++ | 0 | ✗ | 0 | 0 | ✗ | 0 | 0 | - | 0 |
| Histone methylations | H3K4me1 | ✓ | + | + | ✗ | + | ++ | ✓ | + | ++ | - | - |
|  | H3K4me2 | ✓ | +++ | + | ✗ | 0 | 0 | ✓ | + | + | - | 0 |
|  | H3K4me3 | ✓ | +++ | ++ | ✗ | 0 | + | ✗ | ++ | ++ | -- | 0 |
|  | H3K9me3 | ✗ | 0 | 0 | ✗ | - | + | ✓ | - | + | 0 | 0 |
|  | H3K23me2 | ✓ | ++ | NA | ✗ | 0 | NA | ✗ | 0 | NA | 0 | NA |
|  | H3K27me3 | ✓ | -- | 0 | ✗ | -- | 0 | ✓ | --- | + | 0 | - |
|  | H3K36me3 | ✗ | +++ | ++ | ✓ | +++ | +++ | ✓ | +++ | +++ | 0 | 0 |
|  | H3K79me1 | ✓ | ++ | 0 | ✗ | ++ | ++ | ✓ | ++ | ++ | 0 | 0 |
|  | H3K79me2 | ✓ | ++ | 0 | ✗ | + | + | ✓ | ++ | + | 0 | 0 |
|  | H4K20me1 | ✓ | + | NA | ✗ | + | NA | ✓ | + | NA | 0 | NA |
| Histone acetylations | H2AK5ac | ✓ | + | + | ✗ | 0 | 0 | ✗ | 0 | + | 0 | 0 |
|  | H2BK5ac | ✓ | ++ | ++ | ✗ | 0 | + | ✗ | 0 | + | 0 | 0 |
|  | H2BK12ac | ✓ | + | + | ✗ | 0 | + | ✓ | 0 | + | 0 | 0 |
|  | H2BK15ac | ✓ | + | 0 | ✗ | 0 | 0 | ✓ | - | 0 | 0 | 0 |
|  | H2BK20ac | ✗ | 0 | NA | ✗ | 0 | NA | ✗ | 0 | NA | 0 | NA |
|  | H2BK120ac | ✓ | + | + | ✗ | 0 | + | ✗ | 0 | + | 0 | 0 |
|  | H3K4ac | ✓ | + | ++ | ✗ | 0 | + | ✗ | 0 | + | - | 0 |
|  | H3K9ac | ✓ | +++ | ++ | ✗ | 0 | + | ✗ | + | 0 | - | - |
|  | H3K14ac | ✓ | + | + | ✗ | 0 | 0 | ✗ | 0 | 0 | 0 | 0 |
|  | H3K18ac | ✓ | + | ++ | ✗ | 0 | + | ✗ | 0 | + | - | 0 |
|  | H3K23ac | ✓ | + | + | ✗ | 0 | 0 | ✗ | 0 | + | 0 | 0 |
|  | H3K27ac | ✓ | +++ | ++ | ✗ | ++ | ++ | ✗ | ++ | ++ | -- | 0 |
|  | H3K56ac | ✓ | + | NA | ✗ | 0 | NA | ✗ | 0 | NA | 0 | NA |
|  | H4K5ac | ✓ | ++ | NA | ✗ | 0 | NA | ✗ | 0 | NA | 0 | NA |
|  | H4K8ac | ✓ | + | + | ✗ | + | + | ✗ | + | + | - | - |
|  | H4K12ac | ✓ | + | NA | ✗ | + | NA | ✗ | + | NA | - | NA |
|  | H4K91ac | ✓ | + | + | ✗ | 0 | + | ✗ | 0 | 0 | - | 0 |

We obtained RNA sequencing data for about 30 cell or tissue types from the Roadmap data portal ^2^. Gene and exon normalised expression (Transcript per millions, TPM) were calculated by pseudo-mapping of the reads to the human transcriptome using Salmon ^34^ in each sample. Moreover, for every middle exon in each cell type, exon inclusion ratio (or *Ψ*) was calculated (see methods), ranging from 0 to 1 (0 for exons not included, and 1 for exons included in all the transcripts).

The genome-wide histone modification and DNAseI profiles for the cell types corresponding to the transcriptome data were obtained from the Roadmap Epigenomics consortium. For the Whole Genome Bisulfite Sequencing (WGBS) data, we computed three new tracks (figure S1^†^): CpG nucleotide density (consistent across cell and tissue types), CpG methylation ratio (average ratio of methylation at CpG sites in the window), CpG methylation density (number of methylated CpG sites in each window), for each sample, using the WGBS CpG coverage track as a control. For each pair of transcription feature and epigenetic modification, the associations were explored at two levels. (1) Cell or tissue level: for all genes (or exons) in each cell or tissue type and (2) Gene level: for each gene (or exon) across all cell or tissue types. Cell or tissue level analysis allows within-assay comparison of highly and lowly expressed genes (or exons), but is sequence and genomic context (unique for each gene or exon) dependent. Whereas Gene level analysis fixes the sequence and genomic context, but is more sensitive to technical variability across experiments.

The main observations from these analyses are summarised in Table 1 and elaborated in sections below. We have developed a web application to allow exploration of analyses at https://joshiapps.cbu.uib.no/perepigenomics_app/.

### Transcription activity and epigenetic modifications near Transcription Start Sites

Many epigenetic modifications are enriched around the TSS of expressed genes, a region containing gene promoters. To investigate the link between transcription level and epigenetic modifications at the TSS, we generated stack profiles of each epigenetic modification around TSS. When epigenetic modifications were sorted according to gene expression (figure 1A), most histone modifications studied were more abundant at highly expressed genes than at lowly expressed ones. Specifically, only one histone acetylation (H2BK20ac), three histone methylation (H3K9me3, H3K27me3, H3K36me3), and one histone variant H2A.Z did not show a positive correlation with gene expression level. We further noted that only H3K27me3 was more abundant at the TSSs of lowly or not expressed protein coding genes than at the TSSs of highly expressed protein coding genes. H3K27me3 was not present at the TSSs of not expressed, non-protein coding genes in any of the cell or tissue type, highlighting the fact that the associations between an epigenetic modification and transcriptional level may be genetype specific.

**Fig. 1.**
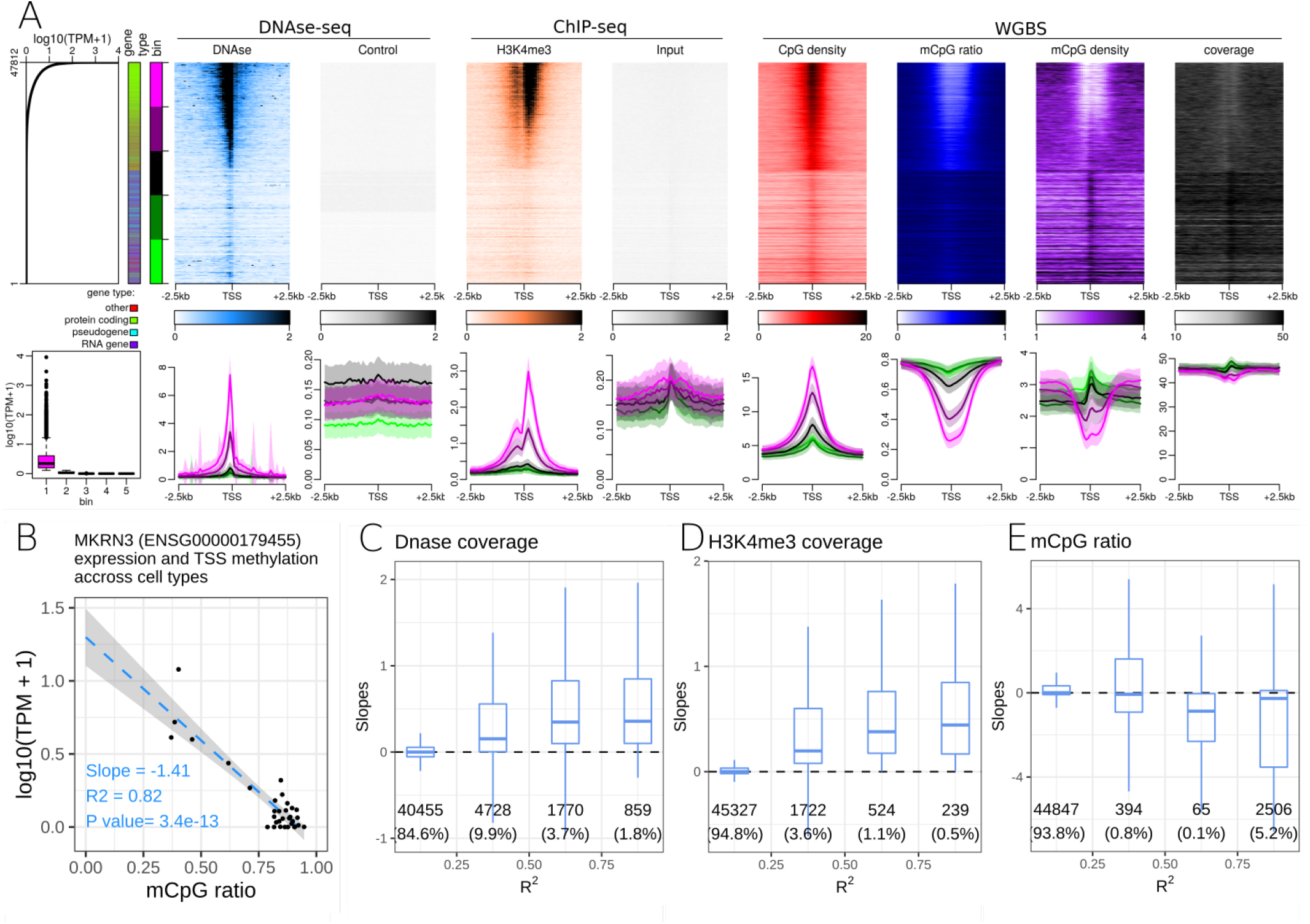
Relationships between epigenetic modifications near the Transcription Start Sites (TSS) and gene expression level. A. Association between epigenetic modifications near the TSS and gene expression level in the small intestine. Upper part, from left to right: Gene expression level in all 47,812 autosomal genes annotated by Gencode. First side bar indicates the gene type (green: protein coding genes, blue: pseudogenes, purple: RNA genes, red: other types of genes), the second side bar indicates the genes sorted according to expression level (figure 1A bottom, 5 bins in total, purple: highly expressed genes, green: lowly expressed genes). Stacked profiles of (i) DNAse-seq and respective control, of (ii) H3K4me3 ChIP-seq and respective input control, of (iii) CpG density, mCpG ratio (mCpG/CpG), mCpG density, and the WGBS coverage near TSS, sorted according to the corresponding gene expression level. Bottom part, from left to right: Boxplot of gene expression levels in each of the 5 expression bins defined in the upper part. Average profiles of DNAse-seq and respective control, of H3K4me3 ChIP-seq and respective input control, CpG density, mCpG ratio, mCpG density and WGBS coverage, ± SEM (Standard Error of the Mean) for each bin of promoters. B-E. Association between epigenetic marks near the TSS and gene expression level across cell types. B. Regression of expression level (in log10(TPM+1)) of the MKRN3 gene and the mean DNA methylation ratio at CpG sites 500bp around the TSS of the MKRN3 gene. Each dot corresponds to a cell type. The slope is negative and the correlation coefficient (*R*^2^) is greater than 0.75 for MKRN3. Similar regressions were generated for each gene and epigenetic modification pair. C-E. Distribution of the slopes from the gene regressions (as in B.) according to the r-squared correlation coefficients for DNAse-seq signal (C.), H3K4me3 ChIP-seq signal (D.) and mCpG ratio (E.) near the TSS of the corresponding genes. Numbers of genes and percentage of genes in each category are displayed below each box.

We explored the trends in peak shapes and noted that a ‘double hill’ (or ‘M’ shape) with a gap at the exact location of the TSS was the most common shape. The ‘gap’ of ChIP-seq signal between the ‘hills’ was located exactly at a sharp peak of DNAseI signal around TSS (figure 1A), denoting a very high DNA accessibility at the TSS. This suggests presence of a nucleosome-free region at promoter terminating at the TSS, with a first nucleosome positioned near the +1 of transcription. This double hill pattern was either symmetric or asymmetric depending on the mark and cell or tissue type (e.g. Stronger peak at the downstream hill than the upstream hill in H3K4me3 in the small intestine, figure 1A). H3K79me2 was particularly strongly asymmetric (figure S2^†^), with a strong peak at around ± 500bp downstream of the TSS across 2 of the 3 cell lines for which the mark was studied in Roadmap Epigenomics.

CpG density around the TSS was positively correlated with gene expression in all cell and tissues in the dataset. CpG methylation ratio and CpG methylation density near gene TSS were negatively correlated with gene expression levels. While CpG methylation ratio showed a flat profile near the TSS of non-expressed genes, CpG methylation density transitioned from a gap at highly expressed genes to a peak at non expressed genes at the TSS (figure 1A).

We further explored these trends for individual gene types. As protein-coding genes formed the majority of all genes, the above observations for all gene types were preserved when the analyses were restricted to protein coding genes only. Separating genes according to gene type highlighted differences in epigenetic profiles at lincRNAs and unprocessed pseudogenes compared to protein coding genes. Processed pseudogenes often had a different relationship between their expression level and the epigenetic status of their promoter. For example, expressed processed pseudogenes showed neither DNAse1 accessibility peak at the TSS, nor an enrichment of active epigenetic modifications at the TSS. While CpG density near the TSS was correlated with processed pseudogene expression level, their promoters did not show any decrease in DNA methylation ratio, resulting in a positive relation between DNA methylation density and processed pseudogene expression. Expressed genes of the ‘antisense’ gene type often mirrored the epigenetic signature of the ‘protein coding’ gene type. For example, H3K79me2 peak was pronounced 500 bp before the TSS of ‘antisense’ genes, whereas it peaked 500bp after the TSS of protein coding genes. However, it is likely that the observed epigenetic signal at antisense genes might be due to the corresponding ‘sense’ gene.

Altogether, gene expression level was positively or negatively correlated with many epigenetic modifications at gene promoters when comparing expressed and not expressed genes within a cell type. The associations between epigenetic modifications and gene expression in a cell or tissue type are promoter sequence and gene context dependent. For example, the CpG density at the TSS is a good predictor of both the gene expression level and the CpG methylation ratio and density (figure 1A). The investigation of associations across cell types is therefore important. To study association between epigenetic modifications and transcription across different cell and tissue types, we calculated linear regression between each epigenetic modification and gene expression level across cell or tissue types. Specifically, the epigenetic signals in ±500bp window around the TSS and the gene expression level (*log*10(*TPM* + 1)) for each gene (figure 1B) were linearly regressed to obtain a slope and a linear correlation coefficient (*R*^2^). For example, the linear regression between the *MKRN3* gene expression level and the average CpG methylation ratio near the TSS of the *MKRN3* gene resulted in a slope of −1.41, an *R*^2^ of 0.82, and a *p*-value of 3.4 · 10^*-*13^. For each epigenetic modification, the distribution of slopes across all genes was plotted against their *R*^2^(figure 1C, 1D and 1E). The epigenetic modifications with positive slopes across cell or tissue types were: H3K4me3 (figure 1D), H3K36me3, H2BK5ac, H3K4ac, H3K9ac, H3K18ac, H3K27ac and chromatin accessibility measured by DNAse1 digestion assay (figure 1C). DNA methylation showed a negative slope for the *R*^2^ greater (figure 1E).

### Transcription activity and epigenetic modifications near Transcription Termination Sites

We repeated the analyses described above at transcription termination sites (TTS). We noted that highly expressed genes tend to be longer than not expressed genes. In short genes, is is difficult to distinguish the effect of epigenetic modifications at TSS from that at the TTS. We therefore defined three classes of genes: short (*≤* 1*kb*), long (*>* 3*kb*), and intermediate length genes, to mitigate this gene-length effect.

While many epigenetic modifications showed a peak (or a gap) centred at the gene TSS, only two modifications were enriched at the gene TTS. First, H3K36me3 displayed a broad hill-shape profile, with a peak at TTS (figure 2A). Levels of H3K36me3 at TTS were positively correlated with gene expression level for different genes within a cell type (figure 2A), and also for gene expression level of the same gene across cell type (figure 2C). In some samples, the H3K36me3 profile was slightly asymmetric at TTS, with more signal in the gene body than after the TTS.

**Fig. 2.**
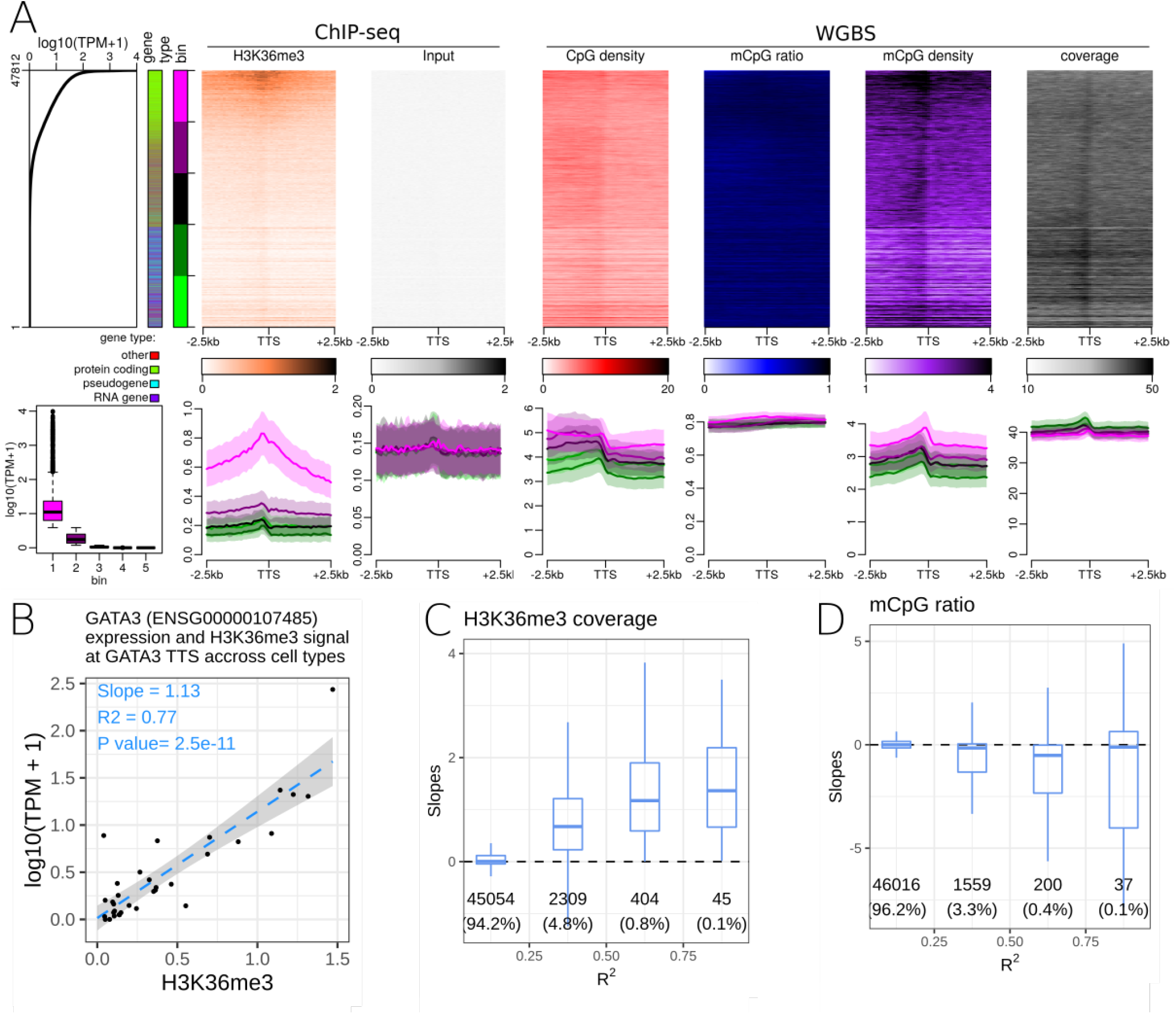
Relationships between epigenentic modifications near the Transcription Termination Sites (TTS) and expression level. A. Association between epigenetic marks near the TTS and gene expression level in the spleen. Upper part, from left to right: Gene expression level in all 47,812 genes annotated by Gencode. First side bar indicates the gene type (green: protein coding genes, blue: pseudogenes, purple: RNA genes, red: other types of genes), the second side bar indicates the 5 bins used in the figure 2A bottom panels (purple: highly expressed genes, green: lowly expressed genes). Stacked profiles of (i) H3K36me3 ChIP-seq and respective input control, of (ii) CpG density, mCpG ratio (mCpG/CpG), mCpG density, and the WGBS coverage near TTS, sorted according to the corresponding gene expression level. Bottom part, from left to right: Boxplot of gene expression level in each of the 5 bins defined in the upper part. Then, average profiles of DNAse-seq and respective control, of H3K4me3 ChIP-seq and respective input control, CpG density, mCpG ratio, mCpG density and WGBS coverage ± SEM (Standard Error of the Mean) for each bin of promoters. B-D. Association between epigenetic marks near the TTS and gene expression level across cell types. B. Regression of expression level (in log10(TPM+1)) of the GATA3 gene with the mean H3L36me3 ChIP-seq signal 500bp around the TTS of the GATA3 gene, where each dot corresponds to a cell or tissue type. The slope was positive and the correlation coefficient (*R*^2^) was greater than 0.75 in this case. Similar regressions were conducted for each gene and epigenetic modification pair. C. and D. Distribution of the slopes from the gene regressions (as in B.) according to the r-squared correlation coefficients for H3K36me3 modification (C.), and mCpG ratio (D.) near the TTS of the corresponding genes. Number of genes and percentage of genes in each category are displayed below each box.

DNA methylation density increased at the TTS (figure 2A). As DNA methylation ratio was nearly constant, the increase of DNA methylation density was mostly due to an increase of CpG density at the TTS. DNA methylation density at the TTS was positively correlated with gene expression level when comparing different genes within a cell type. A weak negative correlation between DNA methylation and gene expression level at TTS was observed for a subset of genes when comparing the same gene across cell and tissue types (figure 2D). This is in agreement with recent findings ^32^. The negative correlation between DNA methylation and gene expression was evident only in ‘long’ genes, and in protein coding genes.

Though most epigenetic modifications did not show enrichment at TTS, four epigenetic modifications (DNAseI accessibility, H3K4me1, H3K79me1, H3K27ac), were positively correlated with expression level of a gene across cell and tissue types. These epigenetic modifications showed no enrichment at TTS, yet the change in expression level was associated with the change in epigenetic modification strength. These modifications could be reflecting broader chromatin organisational features such as topologically associating domain and A/B chromatin domains ^35^.

### Exon transcription and epigenetic modifications at middle exons

Several studies have found a correlation between DNA methylation ^26,27,36^ or histone modifications ^23,30,31^ and exon and splicing events, in either single or a few cell and tissue contexts. We explored whether these observations hold true in the Roadmap Epigenomics dataset. We focused on middle exons and excluded first and last exons of protein coding genes. A total of 16,811 middle exons were expressed in at least one cell or tissue type in the Roadmap dataset. The expression level of an exon was defined as the sum of the TPM of the transcripts including that exon. Similar to TSS and TTS analysis, we performed epigenetic modifications enrichment analysis at middle exons. H3K36me3 showed enrichment at middle exons, correlated with the exon expression level within a cell or tissue type (figure 3A). Changes in H3K36me3 at exons was also strongly associated with exons expression level across cell and tissue types (figure 3D). Though some other epigenetic modifications showed a weak enrichment at exons, similar to input samples used as negative controls, therefore likely reflecting a technical artefact. Epigenetic modifications including H3K4me1, H3K4me3 (figure 3C), H3K27ac, H3K79me1, H3K79me2, H3K9ac, and H3K8ac, were correlated with exon expression level across cell and tissue types, but did not show any enrichment at the middle exons. The H3K4me3 signal, peaked at the gene TSS, and terminated around the start of the first internal exon (figure 3A), resulting in a transition from H3K4me3 marked chromatin to H3K36me3 marked chromatin near the beginning of the second exon of genes.

**Fig. 3.**
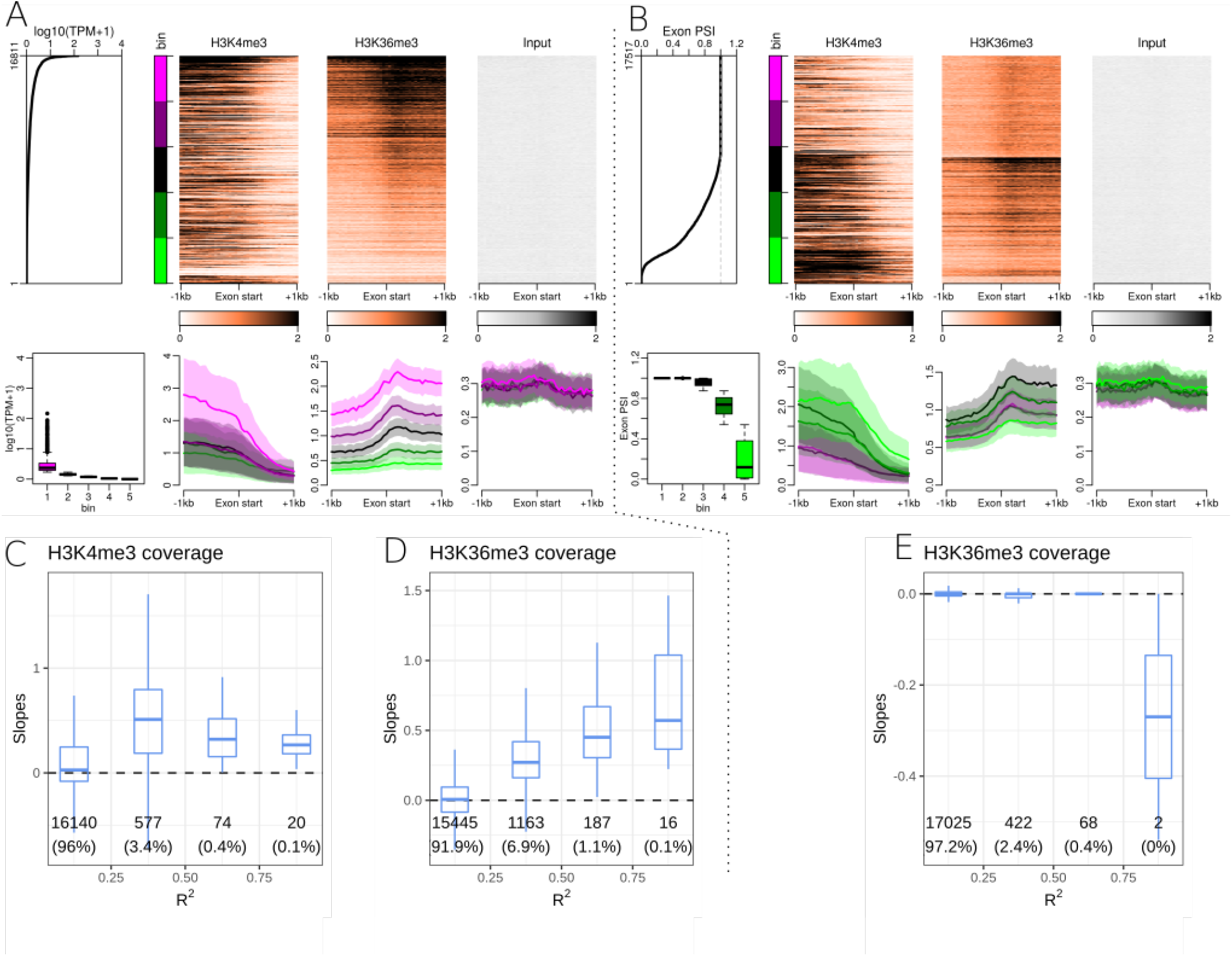
Relationships between epigenentic modifications near middle exons starts and exon expression level or exon inclusion ratio. A. Association between epigenetic marks near middle exons start sites and exon expression levels in the HUES64 cell line. Upper part, from left to right: middle exon expression levels in 16,811 middle exons annotated by Gencode. The side bar indicates 5 bins used in the figure 3A bottom panels (purple: highly expressed exons, green: lowly expressed exons). Then: Stacked profiles of H3K4me3 ChIP-seq, H3K36me3, and input control, sorted according to the exon expression levels. Bottom part, from left to right: Boxplot of exon expression levels in each of the 5 bins defined in the upper part. Then, average profiles of H3K4me3 ChIP-seq, H3K36me3, and respective input control ± SEM (Standard Error of the Mean) for each bin of exons. B. Association between epigenetic marks near middle exons start sites and exon inclusion ratios in the HUES64 cell line. Upper part, from left to right: middle exon inclusion ratios in 17,517 middle exons annotated by Gencode. The side bar indicates 5 bins used in the figure 3B bottom panels (purple: included exons, green: excluded expressed exons). Then: Stacked profiles of H3K4me3 ChIP-seq, H3K36me3, and respective input control, sorted according to the corresponding exon inclusion ratio. Bottom part, from left to right: Boxplot of exon inclusion ratios in each of the 5 bins defined in the upper part. Then, average profiles of H3K4me3 ChIP-seq, H3K36me3, and respective input control ± SEM (Standard Error of the Mean) for each bin of exons. C. and D. Distribution of the slopes from exon expression level regressions and epigenetic mark presence at exon start (± 100 bp) according to the *R*^2^ correlation coefficients for H3K4me3 signal (C.), and H3K36me3 (D.) near the start of the corresponding middle exons. E. Distribution of the slopes from exon inclusion ratio regressions and epigenetic modification at exon start (± 100 bp) grouped by the *R*^2^ correlation coefficients for H3K36me3 modification. Number of exons and percentage of exons in each category are displayed below each box.

There was no difference between DNA methylation ratio at exons and introns, but exons had overall more CpG sites than introns, resulting in an higher DNA methylation density at middle exons than at surrounding introns. DNAse1 accessibility further decreased at exon start (Figure S3^†^), from the already low surrounding accessibility, suggesting that the splicing acceptor site has even lower accessibility than surrounding regions.

### Exon inclusion ratio and epigenetic modifications at middle exons

Exon expression level in a cell type consists of both constitutive and alternative splicing events. To study association between epigenetic modifications and alternate splicing events, we calculated exon inclusion ratio for each exon. The exon inclusion ration was calculated for all transcripts of a given gene (see methods), in each cell or tissue type and sorted using the inclusion ratio (also known as *Psi*). Accordingly, we obtained *Psi* values for 17,517 exons, including 706 exons that were never included in the Roadmap datasets, but were part of genes that were expressed in this dataset. We checked whether epigenetic modifications were correlated with exon inclusion ratio and noted that only a few modifications; namely H3K27ac, H3K4me3, and H3K36me3 were associated with exon inclusion ratio within a cell type (figure 3B). DNA methylation was also associated with exon inclusion ratio. No epigenetic modification showed a significant association with the changes in inclusion ratio at the alternatively included exons (figure 3E, Table 1). There were neverthe-less very weak associations for mCpG ratio, DNAse1, H3K4me1, H3K27me3, H3k9ac and H4K8ac.

### A linear model for gene expression

So far, we analysed the associations between epigenetic modifications and transcription control in a pair-wise manner. In order to better model the combinatorial effect of modifications, we selected 6 epigenetic modifications (DNA methylation, H3K4me1, H3K4me3, H3K9me3, H3K27me3 and H3K36me3) for which epigenetic and transcriptome data was available for 27 cell types in the Roadmap dataset, and regressed a linear model at four transcriptional features: (i) epigenetic modifications around TSS and gene expression level (figure 4A), (ii) epigenetic modifications around TTS and gene expression level (figure 4B), (iii) epigenetic modifications around the start of middle exons and exon expression level (figure 4C), and (iv) epigenetic modifications around the start of middle exons and exon inclusion ratio (figure 4D).

**Fig. 4.**
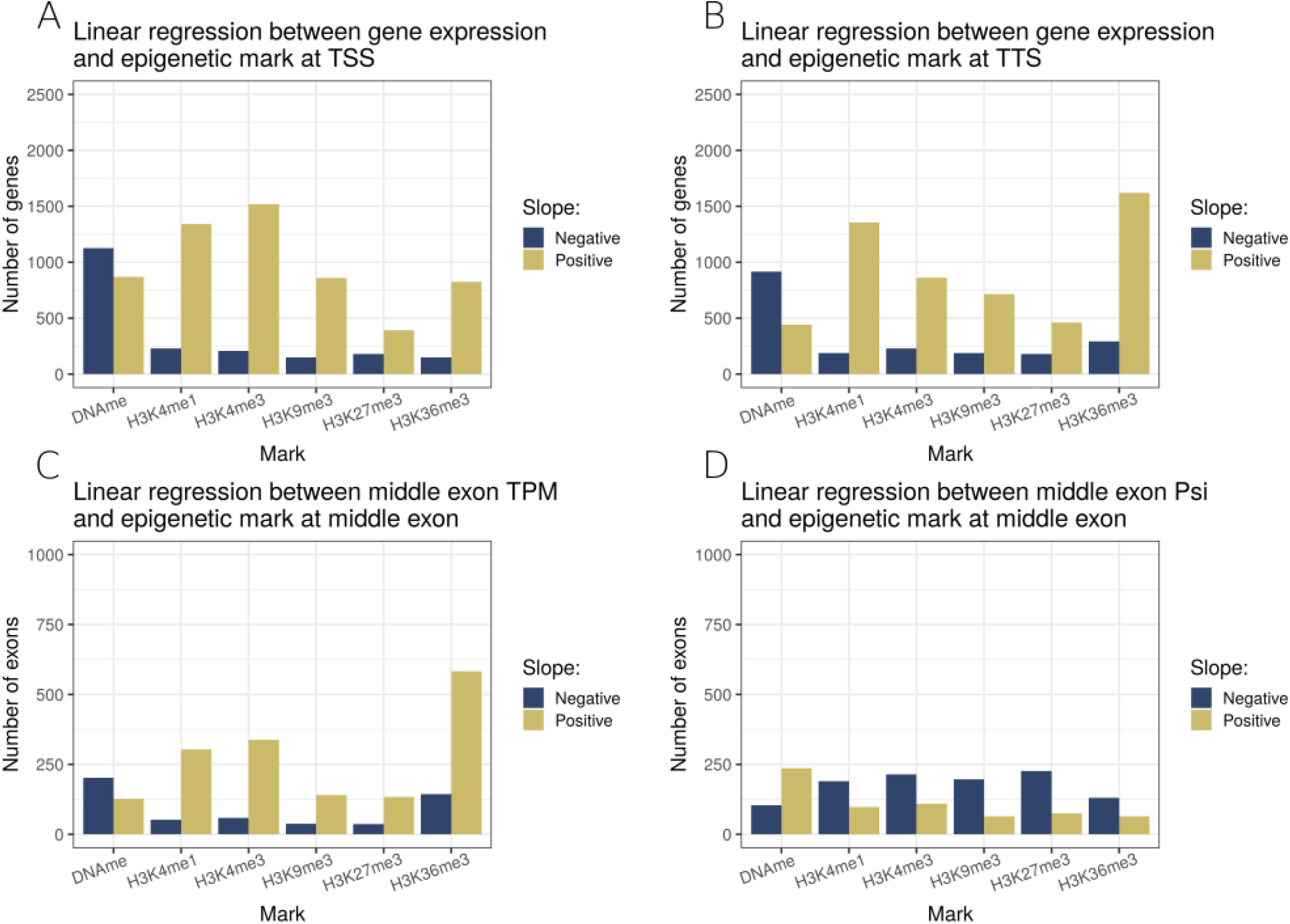
Linear regression models including six epigenetic modifications characterised in 27 cell types. Each bar represents the number of genes or exons with a statistically significant slope (p *<*= 0.01), either positive (golden) or negative (deep blue). A. Linear regression model of gene expression level and the levels of 6 epigenetic modifications near their respective TSS (±500 bp). B. Linear regression model of gene expression level and the levels of 6 epigenetic modifications near their respective TTS (±500 bp). C. Linear regression model of middle exon expression level and the levels of 6 epigenetic modifications near their respective start (±100 bp). D. Linear regression model of middle exon inclusion ratio and the levels of 6 epigenetic modifications near their respective start (±100 bp).

At TSS, amongst the 6 marks studied, DNA methylation was associated with gene repression, while all studied histone methylations were associated with activation of expression level (figure 4A). At TTS, we noted similar pattern as TSS, with a difference that H3K36me3 at TTS was the modification most strongly associated with an increase in expression level, followed by H3K4me1 as the second most positively associated (figure 4B). We noted that restricting the analysis to long genes only, TTS associations with epigenetic modifications were much weaker (figure S4^†^), highlighting that a important portion of the TTS associations were due to short genes for which the promoter marks might confound the TTS signal. H3K36me3 at middle exons was also strongly associated with exon expression levels (figure 4C). Finally, no significant associations were detected for epigenetic modifications (of 6 selected ones) at exon inclusion ratio (figure 4D).

### PEREpigenomics, a web resource to explore associations between epigenetic modifications and transcription features

Using the Roadmap dataset, we unravelled a range of associations between epigenetic modifications and transcription control. We have included a total of 9,024 stacked profiles of epigenetic modifications near TSS, TTS and middle exons, sorted according to the gene expression level, exon expression level or exon inclusion ratio, in about 30 different cell and tissue types. We have also provided stacked profiles for TSS and TTS for each gene type (protein-coding, RNA, pseudogene, other), as well as for short (*≤* 1*kb*), long (*>* 3*kb*) and intermediate length genes. Users furthermore can generate regressions between an epigenetic modification at TSS or TTS and gene expression level across cell and tissue types for a gene of interest, and the same feature is available for middle exons and exon expression level or exon inclusion ratio as well.

We have made the analyses available to users through a web-application at joshiapps.cbu.uib.no/perepigenomics_app/.

## Discussion

In summary, this multi-faceted analysis of the Roadmap Epigenomics data, freely available through a web application, has enabled us confirm known (or previously observed in only one of few cell types) associations between epigenetic modifications and transcription on a large data set and further formulate new hypotheses. We specifically discuss epigenetic associations of previously under-explored transcription features below grouped according to the epigenetic modification.

### Histone modifications

For 28 studied histone modifications near the TSS, 25 were positively correlated with gene expression across genes within a cell type, and a subset of them (16) were also correlated with gene expression when comparing the same gene across cell and tissue types (table 1). H3K27me3 was the only mark that was more abundant at TSS of lowly or not expressed genes than at highly expressed protein coding genes. It is important to note that H3K27me3 modification was not present at non-protein coding genes, highlighting that the negative correlation between H3K27me3 and gene expression is a gene type specific association. H3K9me3 has been associated with closed chromatin ^37,38^, and was not enriched at the TSS of non expressed genes in the Roadmap Epigenomics data set. While many active histone marks showed a TSS asymmetry, with higher signal downstream of the TSS than at the upstream of the TSS. H3K79me1 and H3K79me2 were enriched downstream of the TSS of expressed genes (figure S2^†^), suggesting an association with transcription direction. Indeed, H3K79 methylations are catalysed by the DOT1L enzyme during transcription elongation ^39^. It should be noted that their profile was only available in 5 and 3 cell lines respectively, all derived from the embryonic stem cell line H1. H3K79me1 and H3K79me2 asymmetries were less marked in the H1-derived trophoblast, which could reflect either a relevant biological difference or be caused by experimental issues.

Among histone modifications, H3K36me3 displayed a unique profile across all studied transcription features. The H3K36me3 modification is positively correlated with gene expression at the TTS across genes within a cell type, and also when comparing the same gene across cell and tissue types. While largely absent from the TSS region, it is enriched at all middle exons and on the last exon. We noted that gene body H3K36me3 begins at the start of the second exon, where H3K4me3 peak diminishes. We further explored a potential link between exonic H3K36me3 and alternative splicing. Our approach using exon inclusion ratio revealed that any such association was either weak or restricted to only a few exons. Similar observations have been made by others: Xu *et al* ^31^ noted that from about 3000 alternative splicing events, 800 were positively correlated with changes in H3K36me3 and 700 where negatively correlated with changes in H3K36me3. It should be noted that H3K36me3 modified genomic regions (wide peaks) tend to be an order of magnitude larger than an average exon size (1000 pb vs 100 pb). Altogether, though 10 histone modifications showed some enrichment at exons, the associations between epigenetic modifications and changes in exon inclusion ratio were weak and/or limited to a subset of genes.

### DNA methylation

CpG density is highly variable in the human genome, with CpG depleted or CpG poor regions spanning most of the genome. We noted that CpG density at TSS was strongly associated with gene expression across genes within a cell type, where most expressed genes had a CGI centred at their TSS. Accordingly, at TSS, the mCpG ratio, or the average methylation at CpG sites, was negatively associated with gene expression across genes within a cell type. This trend overlaps with the CpG density where most of CpG deserts are heavily methylated, and most CGI are unmethylated ^40^ (figure S1^†^). Increase in mCpG ratio at TSSs was also associated with the down-regulation of gene expression in the gene level analysis. Non-promoter regions were methylated with mCpG ratios around 85%, this ratio dropped to around 30% at the TSSs of the expressed protein-coding genes. We noted that this not the case for expressed pseudogenes, whose TSSs had higher CpG density than surrounding regions, but remained methylated.

mCpG density is the number of methylated sites at a given genomic region, calculated as a product of the CpG density and the average mCpG ratio at a give genomic region. It has been shown that mCpG density, but not mCpG ratio, was the main driver of the binding of DNA methylation readers of the MBD family ^41^. While many publications focus solely on the mCpG ratio as a metric to evaluate DNA methylation, we argue that multiple metrics provide complementary information. For example, while mCpG ratio stays constant across the promoters of repressed genes, mCpG density peaks near the TSS of repressed genes, suggesting that these regions might preferentially recruit repressive DNA binding proteins (e.g. MBP). DNA methylation density also peaks at the TSSs of expressed processed pseudogenes, as their CGIs remain methylated. At exons, mCpG ratio is as high as at introns, but the CpG density is higher at exons than at introns, resulting in an higher mCpG density.

It has been observed that GC rich region might be more difficult to sequence using some Illumina sequencing protocols, resulting in lower coverage at CGI ^42^. We noted this bias in about half of the WGBS samples, where WGBS coverage at TSS was anti-correlated with gene expression across genes within a cell type. Some WGBS samples were less affected by this bias, while a few showed even an inverse trend, with higher coverage at GC rich regions. Luckily these biases leave mCpG ratio and mCpG density profiles largely unaffected, therefore preserving the validity of the analysis.

### DNA accessibility and nucleosome positioning

DNAse1 assay showed a narrow (<100 bp) peak before and at the TSS of expressed genes. This narrow peak of DNA accessibility matched the location of a dip in the bi-modal signal present in many histone modifications positively correlated with expression level (e.g. H3K4me3 and many histone acetylations). These observations suggest that there is a short nucleosome-free region before the TSS of expressed genes, with a nucleosome positioned just after the +1 of transcription. Such nucleosome positioning effect is well described in yeast ^43^ and in mammals ^44^. Intriguingly, middle exon starts and TTS positions appear to be depleted of DNAse1 signal, even more so than the surrounding regions (figure S3^†^). This suggests that middle and last exon starts are particularly inaccessible regions, maybe due to nucleosome positioning ^45^ or the presence of the splicing machinery at acceptor sites.

### Analysis across cell and tissue types

For each gene and middle exon, we correlated the level of epigenetic modification with expression level or exon inclusion ratio across the different cell and tissue types in the Roadmap dataset. When regressing promoter DNA methylation with gene expression levels, 2,506 genes had a linear regression coefficient *R*^2^ *≥* 0.75 (5.2% of all genes), including 1,173 protein coding genes (6.3% of protein coding genes). The average slope was negative, indicating that for an individual gene, higher level of promoter DNA methylation is associated with lower gene expression level. Conversely, 2,485 genes had a linear regression coefficient *R*^2^ *≥* 0.75 (5.2% of all genes) when regressing H3K4me3 levels at the promoter and gene expression level, including 1,367 protein coding genes (7.3% of protein coding genes). The average slope in this case was positive, i.e. higher level of promoter H3K4me3 was associated with higher gene expression level. Nonetheless, most genes or exons did not have a regression coefficient *≥* 0.25. This might be because many genes and exons might not have large enough epigenetic or expression variability in this dataset. Furthermore, ChIP-seq and DNAse-seq peak height can also be biased by the epigenetic modifications (or accessibility) in the dominant fraction of alleles and cells, as well as biased by changes in ChIP efficiency due to hard-to-control experimental variations.

## Conclusions

In summary, we performed a comprehensive analysis to study links between epigenetic modifications and transcription control using the Roadmap Epigenomic data. The Roadmap Epigenomic histone modifications, whole genome bisulfite sequencing, and RNA-seq data, across diverse human cell and tissue types, allowed us to confirm the generality of previously described relations between epigenetic modifications and transcription control in one or few cell types, as well as to generate new hypotheses about the interplay between epigenetic modifications and transcript diversity. Importantly, our analysis focused on previously under-explored features of transcription control including, transcription termination sites, exon-intron boundaries, middle exons and exon inclusion ratio. We have produced thousands of stack profile plots of epigenetic marks around gene features sorted according to gene expression level, exon expression level or exon inclusion ratio, and filterable by gene type. These plots are made freely available through a web application, joshi-apps.cbu.uib.no/perepigenomics_app/. We hope this web application will serve the community as a resource i) to validate known or previously described epigenetic modifications associated with transcription features ii) as well as an interactive tool to allow exploration of the data as a novel hypotheses generator of epigenetic and transcriptional control.

## Methods

### Data retrieving

Gencode human annotation version 29 (main annotation file), were downloaded from theGencode as gff3 files. Reads from RNA-seq data were retrieved from the European Nucleotide Archive using the the Roadmap sample table as a reference. Roadmap Epigenomics whole genome bisulphite sequencing (WGBS) data sets were downloaded bigwig files of fractional methylation and read coverage from the Roadmap Epigenomics the Roadmap data portal.

Histone modifications and DNAse1 data were downloaded as consolidated, not subsampled, tagAlign files from the the Roadmap data portal.

Altogether, we retrieved 27 RNA-seq and corresponding WGBS datasets, 13 DNAse1 profiles (with matching controls), and 242 histone ChIP-seq datasets in 27 human cell lines or tissues (with 27 matching controls).

### RNA sequencing analysis

RNA-sequencing reads were quantified using Salmon ^34^ v12.0 by pseudoalignement to the human reference genome hg38 and annotations v29 provided by Gencode ^33^. We selected the parameters validateMappings, seqBias, gcBias where on, with biasSpeedSamp equal to 5, and libType equals to A. For samples with biological replicates, the median expression value in TPM samples was used for genes and transcripts. For each exon, exon expression level was calculated as the sum of the TPM of the transcripts including this exon. Exon inclusion ratios as computed as the sum of TPM of transcripts including the exon divided by the TPM value of the gene.

We considered each gene uniquely, by selecting a representative TSS (ory TTS) per gene, from all annotated TSSs (or TTSs). We sorted all transcripts of a gene according to their TSS (or TTS) genomic coordinates and selecting the TSS (or TTS) of the transcript at the middle of the list, i.e. the median TSS (or TTS). The list of middle exons was obtained by taking the shortest transcript of each protein coding gene, then selecting genes with 3 or more exons, and excluding the first and last exons of each transcript. The shortest known transcript isoform was used to ensure that the list of middle exons could not contain first or last exons of other isoforms. Most annotated non-protein coding genes were monoexonic, and were excluded from the exonic analyses.

### Epigenetic modifications data processing

For WGBS, three different tracks were generated from the `FractionalMethylation.bigwig` files using bedtools ^46^ and rtracklayer ^47^: number of CpG sites per window, mean DNA methylation ratio per window, density of mCpG sites per window, using windows of 250 base pair width, sliding by 100 base pairs. The WGBS coverage file was processed similarly to produce a fourth track serving as a control. No post-processing was done for DNAse1 and Histone `tagAlign` files.

### Heatmap generation

Gene types were derived from Gencode ^33^, where the pseudogene category contained all genes with a word “pseudogene”, and the “other” type of gene was defined as neither protein coding, nor pseudogene, nor RNA gene. Genes were binned in 5 groups of equal sizes according to their expression values, and exons in 5 groups according to their expression values or exon inclusion ratio.

For genes with multiple Gencode annotations for TSSs (or TTSs), we sorted all TSSs (or TTSs) according to their genomic coordinate (5’ to 3’, taking into account their orientation) and took the TSS (or TTS) of the transcript in the middle of the list. The list of middle exons was obtained using protein coding genes. For genes with several annotated transcripts, the transcript with the smallest length was selected, first and last exons were filtered, and the remaining exons were kept. Stacked profiles of tracks centred at TSS or TTS were generated using a region of ± 2.5kb around the TSS or TTS, using windows of 100 bases every 100 bases. Stacked profiles of tracks centred at middle exons were drawn using a region of ± 1kb around middle exon starts, with windows of 50 bases every 50 bases. For histone modifications and DNAse1 data, and corresponding input controls, coverage was expressed as FPKM values. For CpG density and mCpG density was defined as the number of (methylated) sites per window (250 bp for TSS or TTS, 100 bp for exons). The mCpG ratio rnged from 0 (all the CpG sites in the window fully unmethylated) to 1 (all the CpG sites in window fully methylated), and the coverage was expressed as number of reads covering a region. For each of the five gene (or exon) bins, we displayed the value distribution as boxplots, and the average profiles ± standard error of the mean (SEM) for each bin. Heatmaps were drawn by a custom script using the following packages: seqplots ^48^, Repitools ^49^, GenomicRanges ^50^, rtracklayer ^47^, and plotrix ^51^.

### Regression analysis

At each gene TSS (or TTS), the epigenetic modification level in a sample was averaged in the ± 500 bp region around the TSS (or TTS) and a linear regression was calculated for each gene using expression values in *log*10(*TPM* + 1). Epigenetic marks at the ± 100 bp region around middle exon starts were regressed with either *log*10(*TPM* + 1) of the exon or the exon inclusion ratio. For each regression the slope and regression coefficient (*R*^2^) were obtained, using dplyr ^52^, purrr ^53^, and broom ^54^.

### Web application and script availability

PEREpigenomics is developed in Shiny ^55^. Source code and data of the application are available at forgemia.inra.fr/guillaume.devailly/perepigenomics_app. Scripts used to process the data and generate the plots can be found at: github.com/gdevailly/perepigenomicsAnalysis.

## Author’s contributions

GD and AJ designed the analyses and the web applications, and wrote this manuscript. GD performed the analysis and developed the web application.

## Acknowledgements

GD was funded for this work by the People Program (Marie Curie Actions FP7/2007-2013) under REA grant agreement No PCOFUND-GA-2012-600181. AJ is supported by the Bergen Research Foundation Grant no. BFS2017TMT01. The authors would like to thanks Olaf Sarnow, Kjell Petersen and Stanislav Oltu for their help with the web server configuration, and Anna Mantsoki and Barry Horne for her help at the beginning of the project.

## Abreviations

TSS: Transcription Start Sites
TTS: Transcription Termination Sites
TPM: Transcript Per Million of reads
CpG: Cytosine - Guanine dinucleotide
CGI: CpG island
*Psi*: Exon inclusion ratio
WGBS: Whole Genome Bisulfite Sequencing
H3K9me3: tri-methylation of lysine 9 of histone 3
H4K5ac: acetylation of lysine 5 of histone 4

## Supplementary figures

**Supplementary Figure S1:**
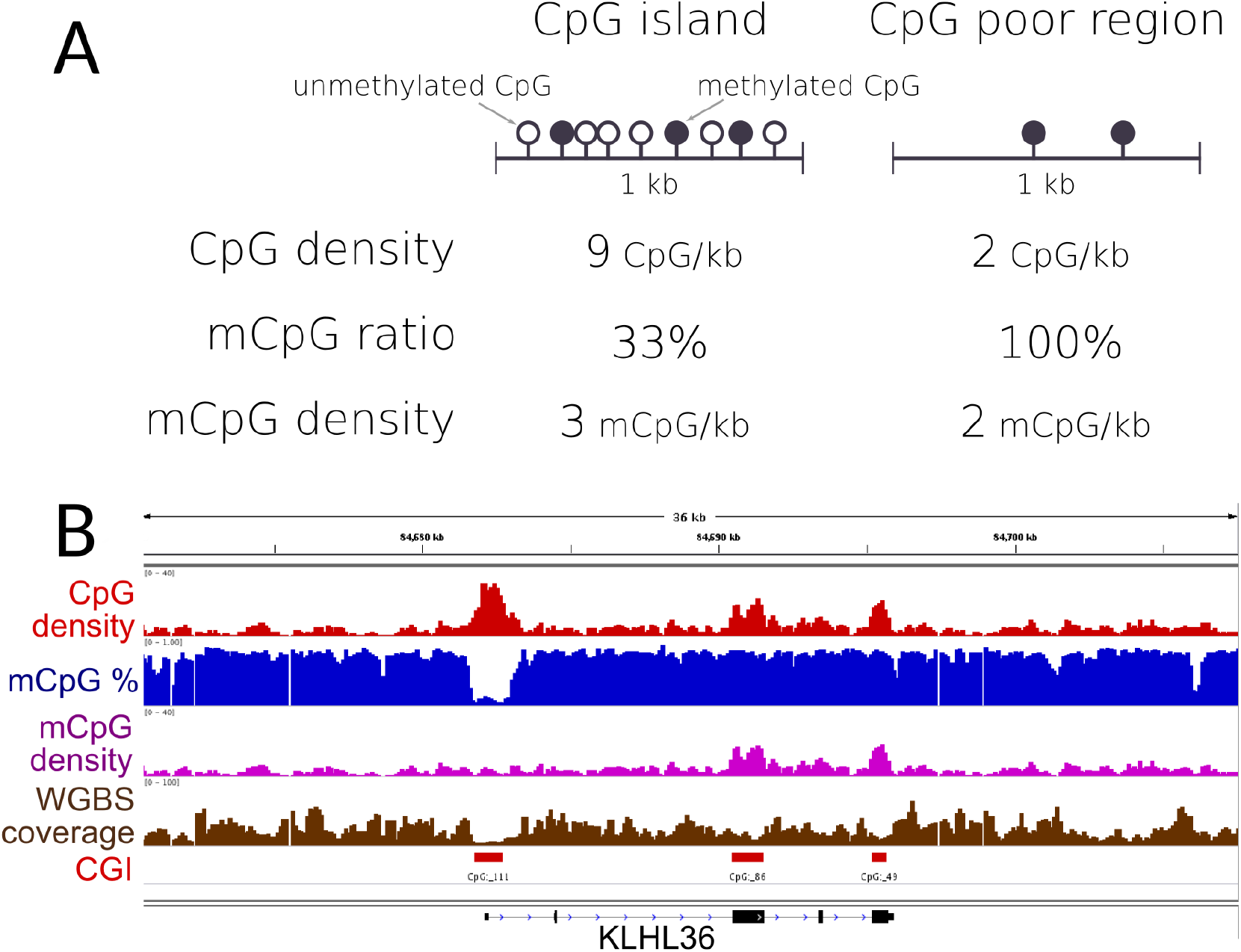
WGBS is analysed through CpG density, mCpG ratio and mCpG density metrics. The mCpG ratio an mCpG density metrics can behave differently if comparing regions with different CpG density. A. On these hypothetical 1 kb genomic windows, the CpG rich region (left) has an lower mCpG ratio than the CpG poor region (right), but an higher mCpG density. B. Four tracks have been obtained from WGBS data (here from the H1-hESC cell line). The 36kb region including the gene KLHL36 is shown. From top to bottom: The CpG density tracks, higher at the three CpG islands in the region. The mCpG ratio, showing high values (> 80%) everywhere but at the first CpG island. The mCpG density, showing low values everywhere but at the second and thris CpG island. Finaly, the WGBS coverage is used as a control track. Here it shows a lower coverage at the three CpG islands.

**Supplementary Figure S2:**
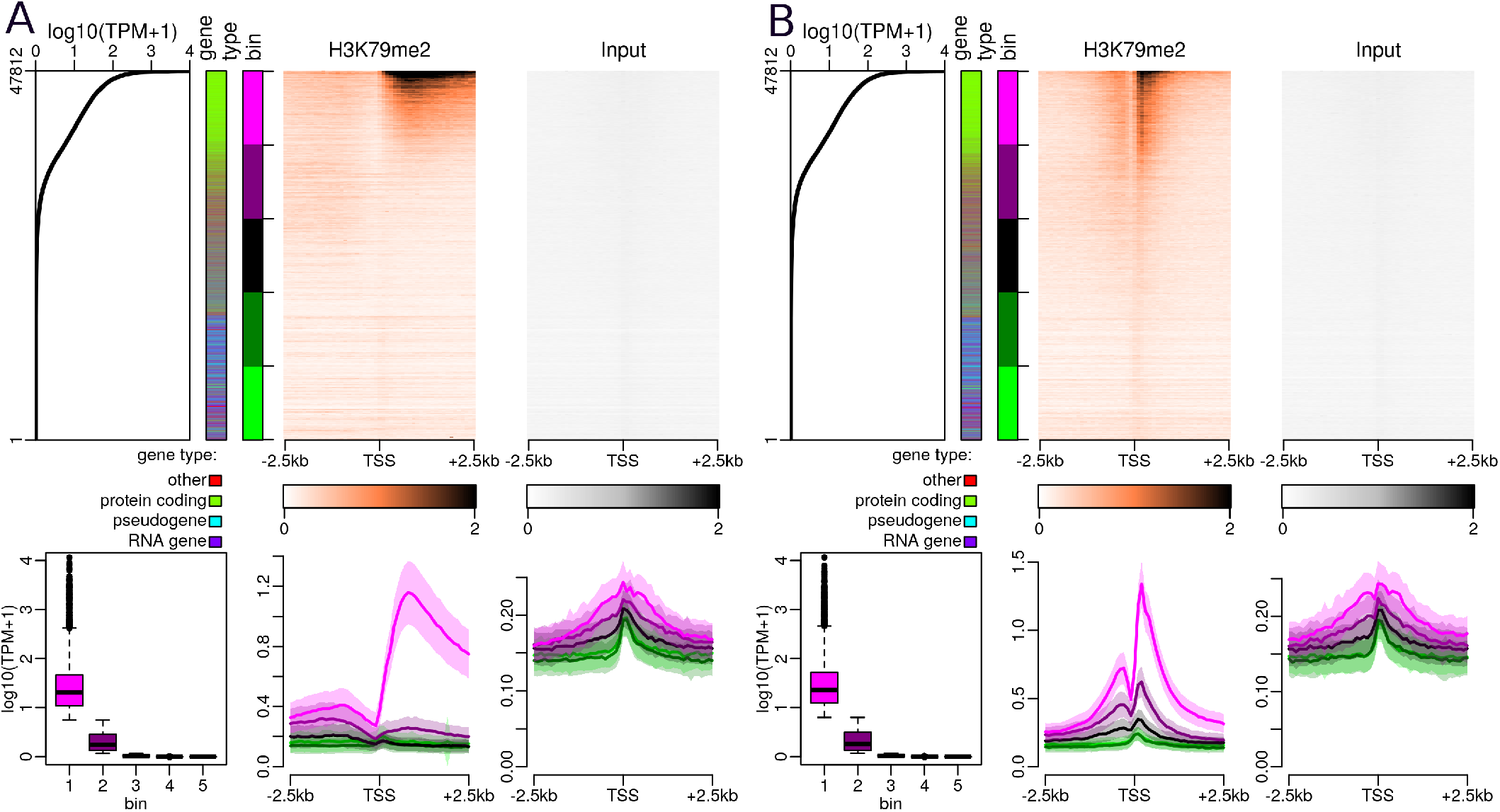
H3K79me2 near the transcription start sites in H1-BMP4 derived mesoderm (A) or H1-BMP4 derived throphoblast (B). Upper parts, from left to right: Gene expression level in all 47,812 autosomal genes annotated by Gencode. First side bar indicates the gene type (green: protein coding genes, blue: pseudogenes, purple: RNA genes, red: other types of genes), the second side bar indicates the genes sorted according to expression level (5 bins in total, purple: highly expressed genes, green: lowly expressed genes). Stacked profiles of H3K79me2 ChIP-seq and respective input control, sorted according to the corresponding gene expression level. Bottom parts, from left to right: Boxplot of gene expression levels in each of the 5 expression bins defined in the upper part. Average profiles of H3K79me2 ChIP-seq and respective input control, ± SEM (Standard Error of the Mean) for each bin of promoters.

**Supplementary Figure S3:**
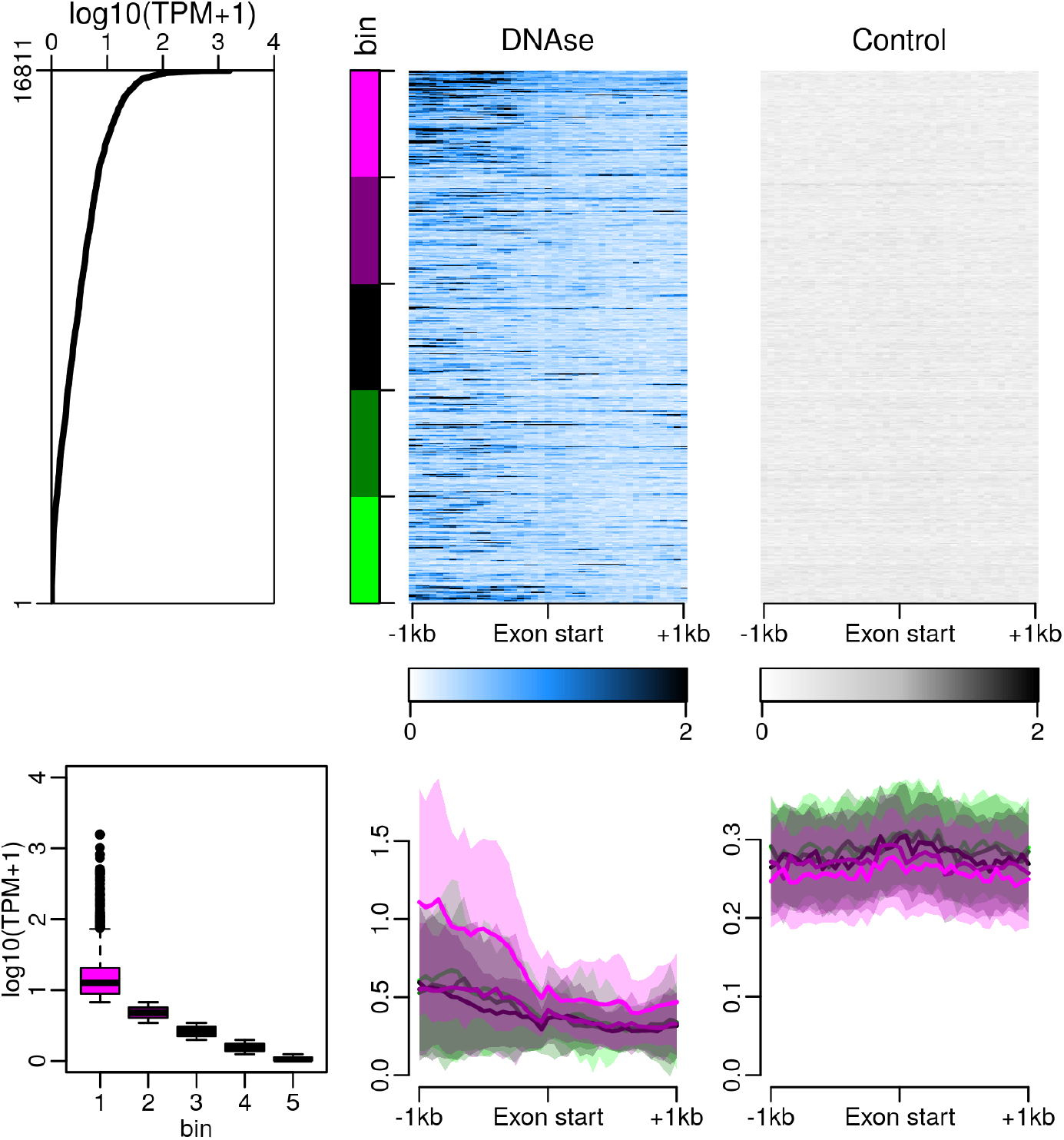
DNAse1 profile around middle exon start sites in pancreas show a narrow decrease in DNAse accessibility near splicing acceptor sites. Upper part, from left to right: middle exon expression levels in 16,811 middle exons. The side bar indicates 5 bins used in the figure S3 bottom panels (purple: highly expressed exons, green: lowly expressed exons). Then: Stacked profiles of DNAse1 profile and control, sorted according to the exon expression levels. Bottom part, from left to right: Boxplot of exon expression levels in each of the 5 bins defined in the upper part. Then, average DNAse profiles and control ± SEM (Standard Error of the Mean) for each bin of exons.

**Supplementary Figure S4:**
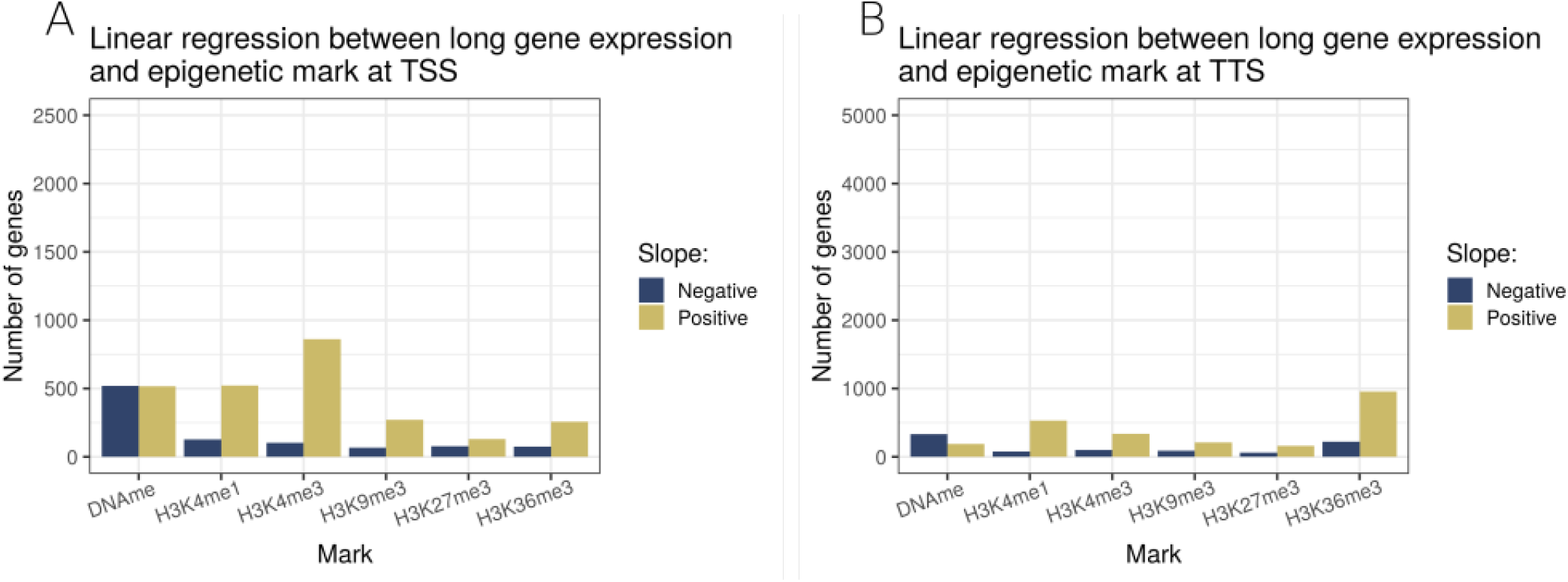
Linear regression models including six epigenetic modifications characterised in 27 cell types, focusing on the 25,068 long genes (> 3kb, amongst 47,812 analysed genes). Each bar represents the number of genes with a statistically significant slope (p *<*= 0.01), either positive (golden) or negative (deep blue). A. Linear regression model of long gene expression level and the levels of 6 epigenetic modifications near their respective TSS (±500 bp). B. Linear regression model of long gene expression level and the levels of 6 epigenetic modifications near their respective TTS (±500 bp).

